# PhyloSuite: an integrated and scalable desktop platform for streamlined molecular sequence data management and evolutionary phylogenetics studies

**DOI:** 10.1101/489088

**Authors:** Dong Zhang, Fangluan Gao, Wen X. Li, Ivan Jakovlić, Hong Zou, Jin Zhang, Gui T. Wang

**Author notes:** **Corresponding author:** (GTW).

## Abstract

Multi-gene and genomic datasets have become commonplace in the field of phylogenetics, but many of the existing tools are not designed for such datasets, which makes the analysis time-consuming and tedious. We therefore present PhyloSuite, a user-friendly workflow desktop platform dedicated to streamlining molecular sequence data management and evolutionary phylogenetics studies. It employs a plugin-based system that integrates a number of useful phylogenetic and bioinformatic tools, thereby streamlining the entire procedure, from data acquisition to phylogenetic tree annotation, with the following features: (i) point-and-click and drag-and-drop graphical user interface, (ii) a workspace to manage and organize molecular sequence data and results of analyses, (iii) GenBank entries extraction and comparative statistics, (iv) a phylogenetic workflow with batch processing capability, (v) elaborate bioinformatic analysis for mitochondrial genomes. The aim of PhyloSuite is to enable researchers to spend more time playing with scientific questions, instead of wasting it on conducting standard analyses. The compiled binary of PhyloSuite is available under the GPL license at https://github.com/dongzhang0725/PhyloSuite/releases, implemented in Python and runs on Windows, Mac OSX and Linux.

## Introduction

Advancements in next-generation sequencing technologies (Metzker, 2009) have resulted in a huge increase in the amount of genetic data available through public databases. While this opens a multitude of research possibilities, retrieving and managing such large amounts of data may be difficult and time-consuming for researchers who are not computer-savvy. A standard analytical procedure for phylogenetic analysis is: selecting and downloading GenBank entries, extracting target genes (for multi-gene datasets, such as organelle genomes) and/or mining other data, sequence alignment, alignment optimization, concatenation of alignments (for multi-gene datasets), selection of best-fit partitioning schemes and evolutionary models, phylogeny reconstruction, and finally visualization and annotation of the phylogram. This can be very time-consuming if different programs have to be employed for different steps, especially as they often have different input file format requirements, and sometimes even require manual file tweaking. Therefore, multifunctional, workflow-type software packages are becoming increasingly needed by a broad range of evolutionary biologists (Smith, 2015). Specifically, as single-gene datasets are rapidly being replaced by multi-gene or genomic datasets as a tool of choice for phylogenetic reconstruction (Degnan and Rosenberg, 2009; Rivera-Rivera and Montoya-Burgos, 2016), automated gene extraction from genomic data and batch manipulation in some of the above steps, like alignment, are becoming a necessity.

Although there are several tools in existence, designed to streamline this process by incorporating some or all of the steps mentioned above, none of these incorporate all of the above functions in a manner suitable for current trends in phylogenetic analyses (see detailed comparison in Supplementary data). Therefore, we present PhyloSuite, a versatile tool designed to incorporate all of the functions described above, including a series of different phylogenetic analysis algorithms, into a single workflow that does not require programming skills, has an intuitive graphical user interface (GUI), workspace, batch mode, extensive plugins support, inbuilt updating function, etc. (Fig. 1). This tool aims to be accessible to all scientists, streamline the phylogenetic analysis procedure, and allow scientists to focus on solving scientific questions rather than waste time on toying with different scientific software programs.

**Fig. 1.**
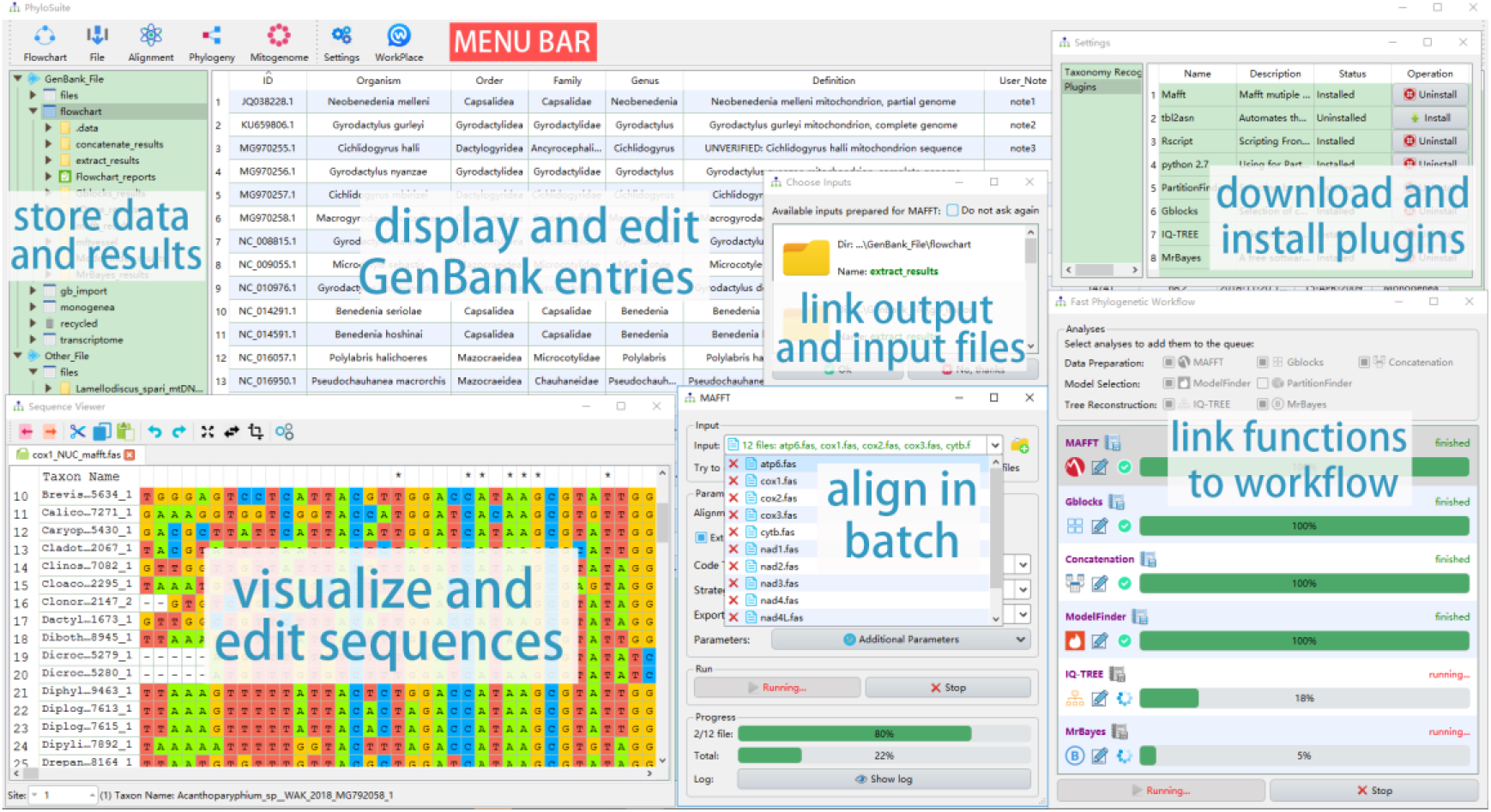
The interface and the main functions of PhyloSuite

## Implementation

PhyloSuite is a user-friendly stand-alone GUI-based software written in Python 3.6.7 and packaged and tested on Windows, Mac OSX and Linux. The functions are (Fig. 1, Supplementary data): (i) retrieving, extracting, organizing and managing molecular sequence data, including GenBank entries, nucleotide and amino acid sequences, and sequences annotated in Word documents; (ii) batch alignment of sequences with MAFFT (Katoh and Standley, 2013), for which we added a codon alignment (translation align) mode; (iii) batch optimization of ambiguously aligned regions using Gblocks (Talavera and Castresana, 2007); (iv) batch conversion of alignment formats (FASTA, PHYLIP, PAML, AXT and NEXUS); (v) concatenation of multiple alignments into a single dataset and preparation of a partition file for downstream analyses; (vi) selection of the best-fit evolutionary model and/or partitioning scheme using ModelFinder (Kalyaanamoorthy, et al., 2017) or PartitionFinder (Lanfear, et al., 2017); (vii) phylogeny reconstruction using IQ-TREE (maximum likelihood) (Nguyen, et al., 2015) and/or MrBayes (Bayesian inference) (Ronquist, et al., 2012); (viii) linking the functions from (ii) to (vii) into a workflow; (ix) annotating phylogenetic trees in the iTOL webtool (Letunic and Bork, 2016) using datasets generated by the (i) function; (x) comprehensive bioinformatic analysis of mitochondrial genomes (mitogenomes); (xi) visualization and editing of sequences using a MEGA-like sequence viewer; (xii) storing, organizing and visualizing data and results of each analysis in the PhyloSuite workspace.

### Genetic data management

PhyloSuite provides a flexible GenBank entries processing function (see Supplementary data). GenBank files can be imported either directly, or via a list of IDs, which PhyloSuite will automatically download from the GenBank. Almost all of the information in the annotation section of a GenBank record can be extracted and displayed in the GUI. Additionally, the information can be standardized in batch using a corresponding function or edited manually in the GUI, ambiguously annotated mitogenomic tRNA genes can be semi-automatically reannotated using ARWEN (Laslett and Canback, 2008), and taxonomic data can be automatically retrieved from WoRMS and NCBI Taxonomy databases. The ‘extract’ function allows users to extract genes in batches, as well as generate an assortment of statistics and dataset files (iTOL datasets). The extracted results can be used for downstream analyses without additional manipulation. The nucleotide and amino acid sequences can be visualized and edited in a MEGA-like explorer equipped with common functions (reverse complement, etc.). Importantly, PhyloSuite can parse the sequence annotations recorded in a Word document via the inbuilt ‘comment’ function, and generate a GenBank file and an *.sqn file for direct submission to the GenBank. This function provides a novel and simple way to annotate genetic sequences, which shall benefit researchers who are not computer-savvy.

### Phylogenetic analysis workflow

By allowing users to combine seven plugin programs/functions and execute them sequentially, PhyloSuite streamlines the evolutionary phylogenetics analysis (see Supplementary data). The standard execution order of these functions is: MAFFT, Gblocks, Concatenation, ModelFinder or PartitionFinder2, MrBayes and/or IQ-TREE. The results of upstream functions are directly prepared as the input for downstream functions, so only the first function of each workflow requires an input file(s). Functions can also be used in a non-standard order and/or separately, in which case PhyloSuite will automatically search for available input files (results of other tools) in the workspace. Before starting the workflow, PhyloSuite will summarize the parameters of each function, allowing the user to check and modify them, or autocorrect conflicting parameters, such as sequence types. Once a workflow is finished, PhyloSuite will describe the settings of each function as well as present the references for each plugin program in the GUI.

### Bioinformatics analysis for mitogenomic data

PhyloSuite was originally designed for, and its major comparative advantages are in, the mitochondrial genomics analyses. There is a specialized configuration available for the extraction of mitogenomic features. In addition to gene extraction, PhyloSuite will generate a dozen of statistics and dataset files useful for downstream analyses (see Supplementary data). The ‘itol’ dataset can be used to annotate the obtained phylogram (colorize lineages, map gene orders, etc.). The gene order file can be used to conduct gene order analysis with CREx (Bernt, et al., 2007) or treeREx (Bernt, et al., 2008). The tables generated include the list of mitogenomes and overall statistics, annotation, nucleotide composition and skewness, relative synonymous codon usage (RSCU) and amino acid usage. The RSCU figure (see Fig. 3 in Zhang, et al. (2017)) can be drawn using the RSCU table and ‘Draw RSCU figure’ function. The annotation table can be used to compare genomic annotations and calculate pairwise similarity of homologous genes with ‘Compare table’ function (see Table 1 in Zhang, et al. (2018)). In the future, PhyloSuite aims to gradually extend these analyses to other small genomes (organelles, viruses, etc.).

## Discussion

PhyloSuite links the management of genetic sequence data and a series of phylogenetic analysis tools, thereby simplifying and speeding up multi-gene based phylogenetic analyses, from data acquisition to phylogram annotation. In summary, highlights of PhyloSuite include: (i) a user-friendly workspace to visualize, organize, manipulate and store sequence data and results; (ii) flexible GenBank entries processing (standardization, reannotation, etc.); (iii) batch data processing capability and workflow; (iv) a state of the art mitogenomic bioinformatics analysis. Although PhyloSuite is designed primarily to allow non-computer-savvy users to drag-and-drop and point-and-click their way through the phylogenetic analysis, experienced scientists will also find it useful to streamline their research, store and manage results, and increase productivity. It will especially benefit evolutionary biologists who wish to test the effects of different datasets and analytical methods on the phylogenetic reconstruction.

## Acknowledgements

The authors would like to thank Dr. Xiao-Qin Xia for modifying the manuscript and Mr. Cheng-En Zheng for technical assistance in the software development.

## Supplementary Information

### Overview

PhyloSuite is designed to address the global trend towards multi-gene based phylogenetic analyses (Degnan and Rosenberg, 2009; Rivera-Rivera and Montoya-Burgos, 2016): this software program incorporates and streamlines all steps included in such analyses, from data acquisition to phylogenetic tree visualization and annotation (Fig. S1).

**Fig. S1.**
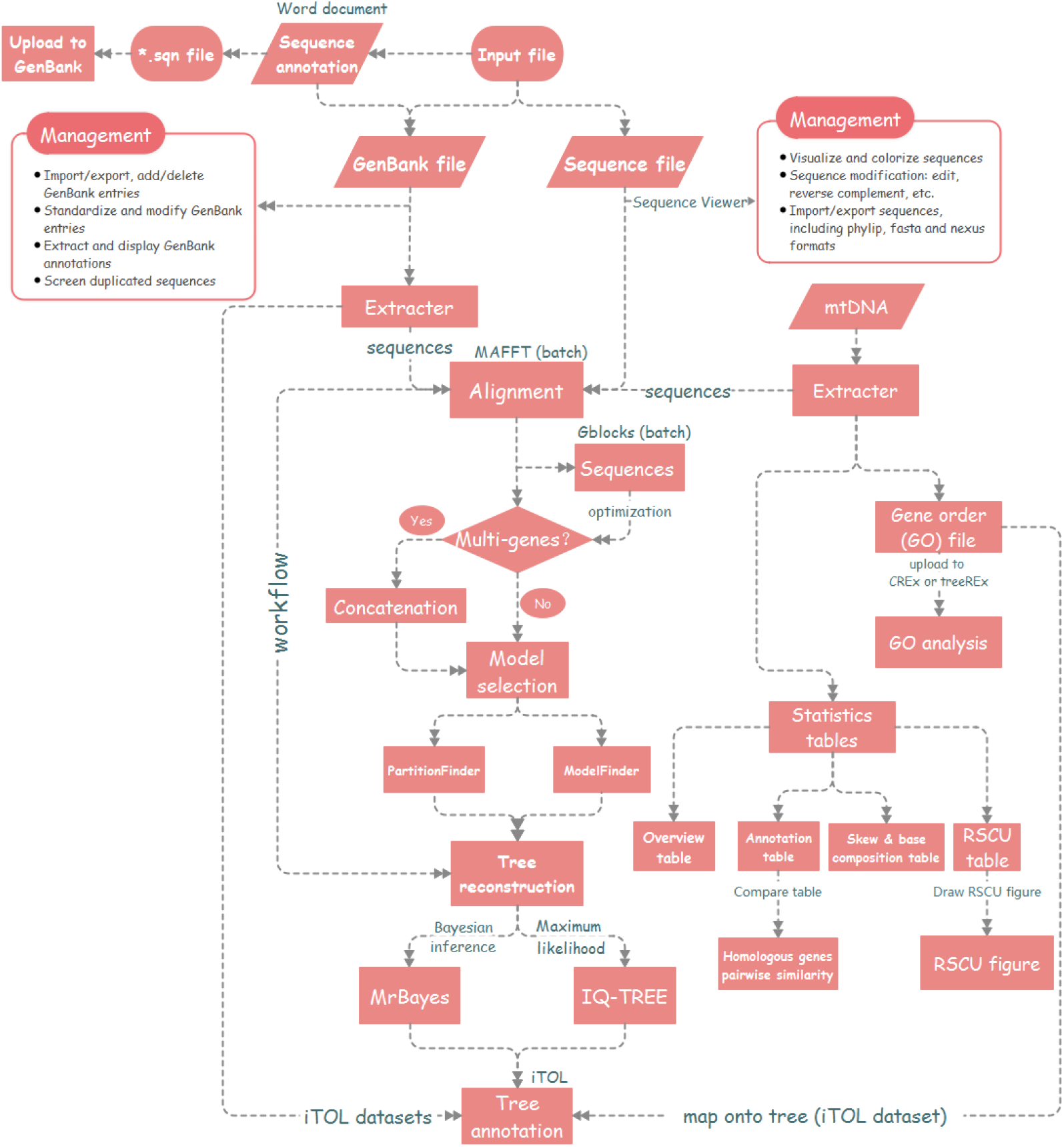
The workflow diagram of PhyloSuite.

### Comparison with extant software programs

Although the extant software programs possess some of the abilities of PhyloSuite, none of them incorporate all functions necessary for a streamlined multi-gene phylogenetic analysis, from data retrieval to the phylogenetic tree annotation (Fig. S3). For example, FeatureExtract (Wernersson, 2005) and Geneious (Kearse, et al., 2012) can extract the annotations from GenBank files, but downstream analysis is not fully automated, so some manual data handling is required, especially for multi-gene datasets. Armadillo (Lord, et al., 2012), EPoS (Griebel, et al., 2008) and MEGA (Kumar, et al., 2016) do not possess the batch processing capability, which is indispensable for multi-gene datasets. Additionally, data partitioning and best-fit partitioning scheme estimation are also pivotal for multi-gene dataset-based phylogenetic analyses (Blair and Murphy, 2011; Lanfear, et al., 2012), but most other software programs lack this function, including Geneious, MEGA, Galaxy Workflow (Oakley, et al., 2014), etc. Although, MEGA and EPoS possess the ability to use the output of one tool directly as the input for another tool, they cannot link several functions into a single run (workflow). Probably the closest to meeting the described requirements is Geneious, but this is a commercial bioinformatics software, so it may not be an ideal option (i.e. too expensive) for all scientists, especially for students.

**Fig. S3.**
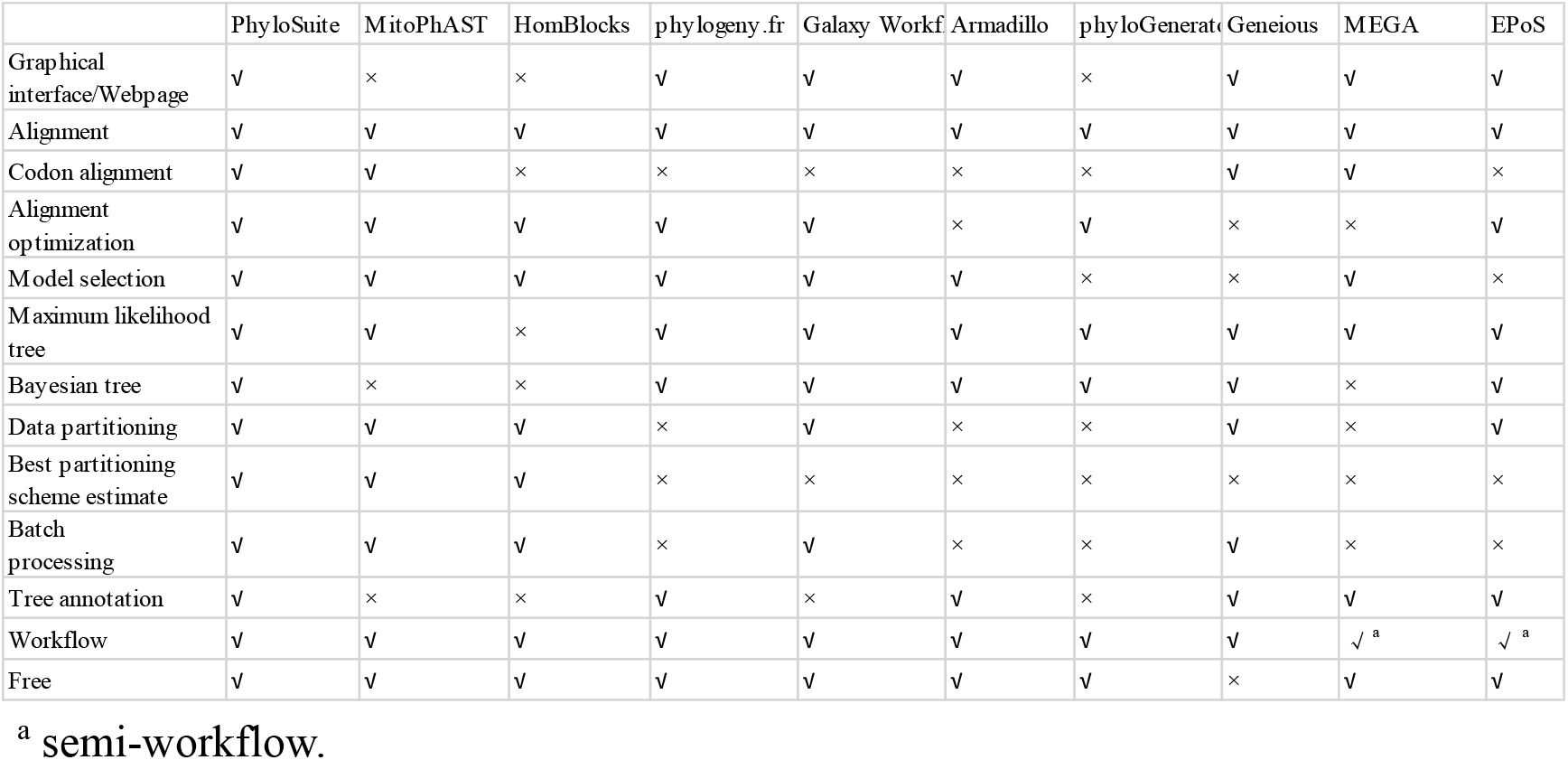
Comparison of PhyloSuite with software programs with similar functions.

### Functions and capabilities

Taking a recently published mitogenomic paper (Zhu, et al., 2018) as an example, using the ‘extract’ function user can quickly conduct most of the analyses reported in that paper, and generate similar tables and figures: (i) mitogenome list and overall statistics table (Fig. S4, Table 1 in Zhu et al.), (ii) annotation table (Fig. S5, Table 2 in Zhu et al.), (iii) nucleotide composition and skewness table (Fig. S6, Table 3 in Zhu et al.), (iv) relative synonymous codon usage (RSCU) table (Fig. S7) and figure (Fig. S8, Fig. 2B in Zhu et al.), (v) amino acid usage statistics file (Fig. S9) used to draw Fig. 2A in Zhu et al., and (vi) reconstruct and annotate (in iTOL) phylogenetic trees (Fig. 5 in Zhu et al.) using the extracted genes. In comparison, most of the tables in that paper were made manually by the author, which is time-consuming, tedious and error-prone. Beyond these, several additional analyses are available: (i) gene order file is generated, which can be used to map gene orders of mitogenomes onto the phylograms in iTOL (Fig. S10, also see Fig. 6 in Zhang, et al. (2018)) and conduct gene order analysis with CREx (Bernt, et al., 2007) or treeREx (Bernt, et al., 2008), (ii) statistics for individual genes, including size, start and terminal codons, base composition and skews (Fig. S11), (iii) general statistics table, which can be used to draw skewness and base content figure (Fig. S12, also see Fig. 1 in Zhang, et al. (2018)), (iv) comparison of genomic annotations and pairwise similarity calculation for homologous genes (Fig. S13, Table 1 in Zhang, et al. (2018).

**Fig. S4.**
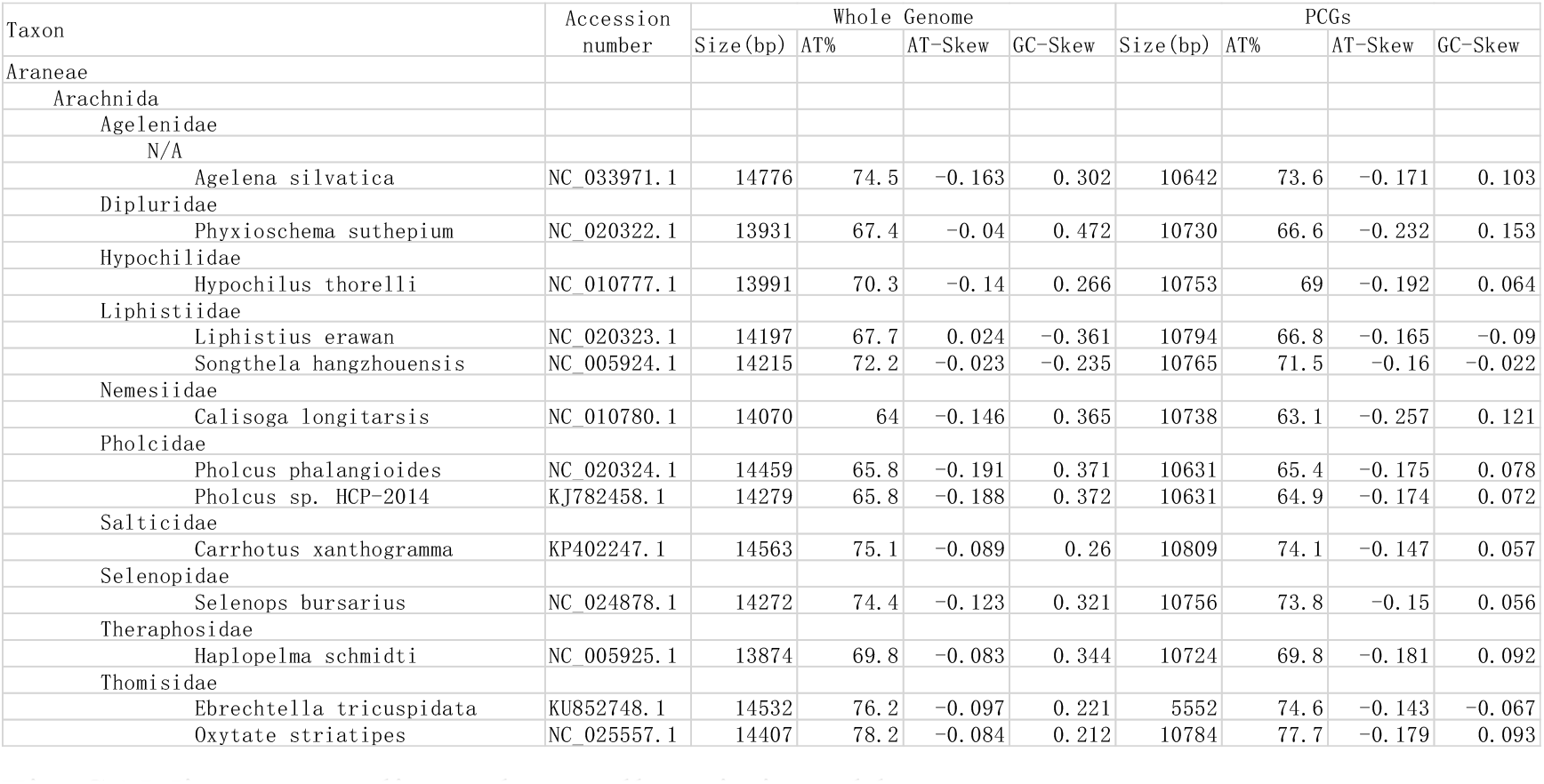
Mitogenome list and overall statistics table.

**Fig. S5.**
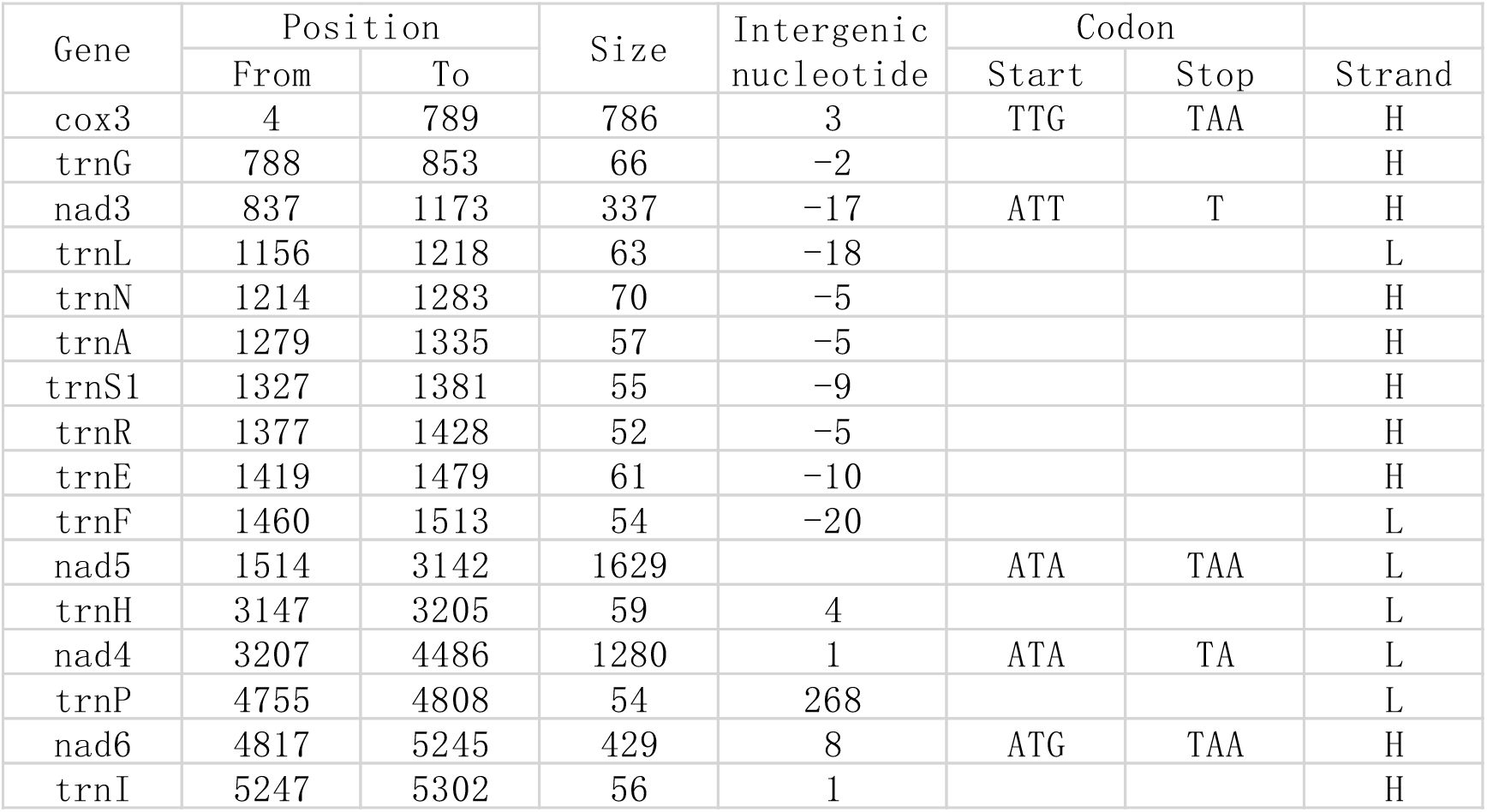
Annotation table.

**Fig. S6.**
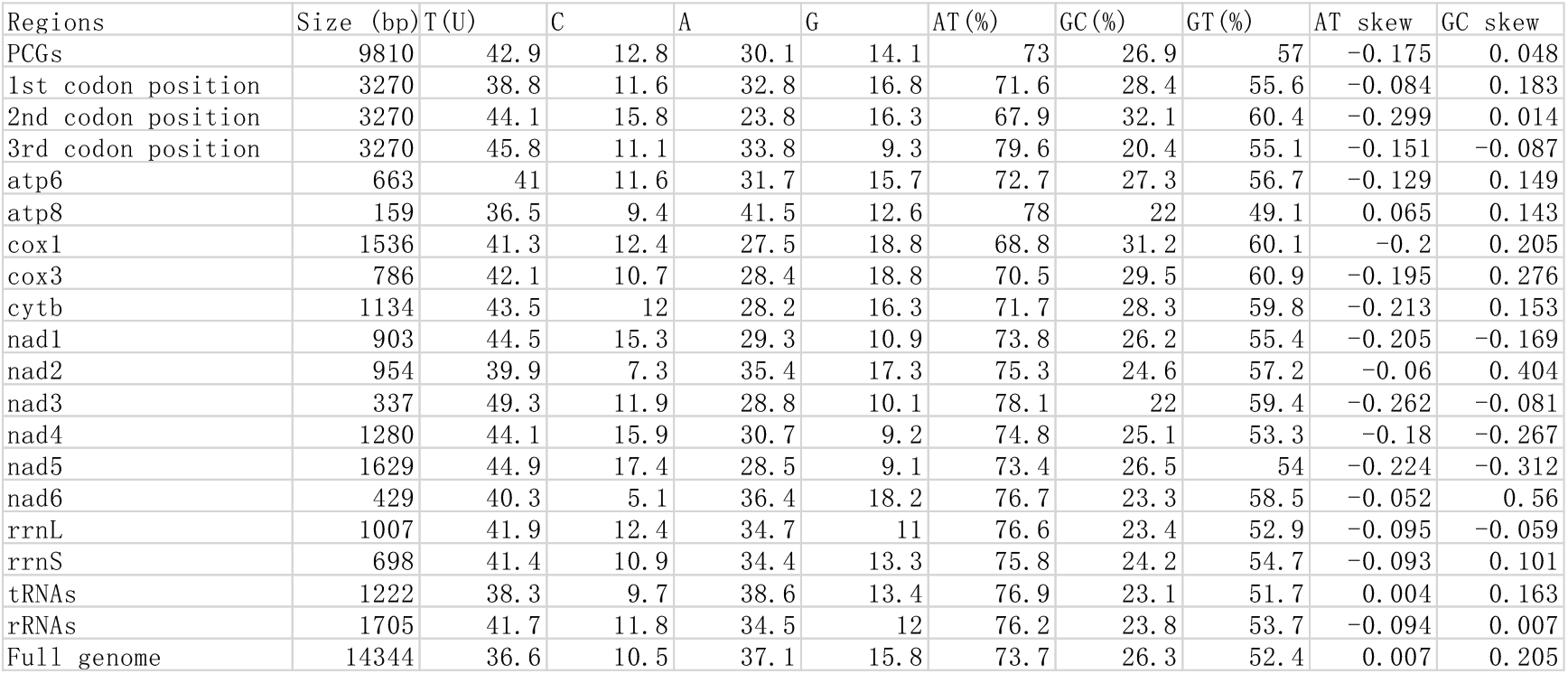
Nucleotide composition and skewness table.

**Fig. S7.**
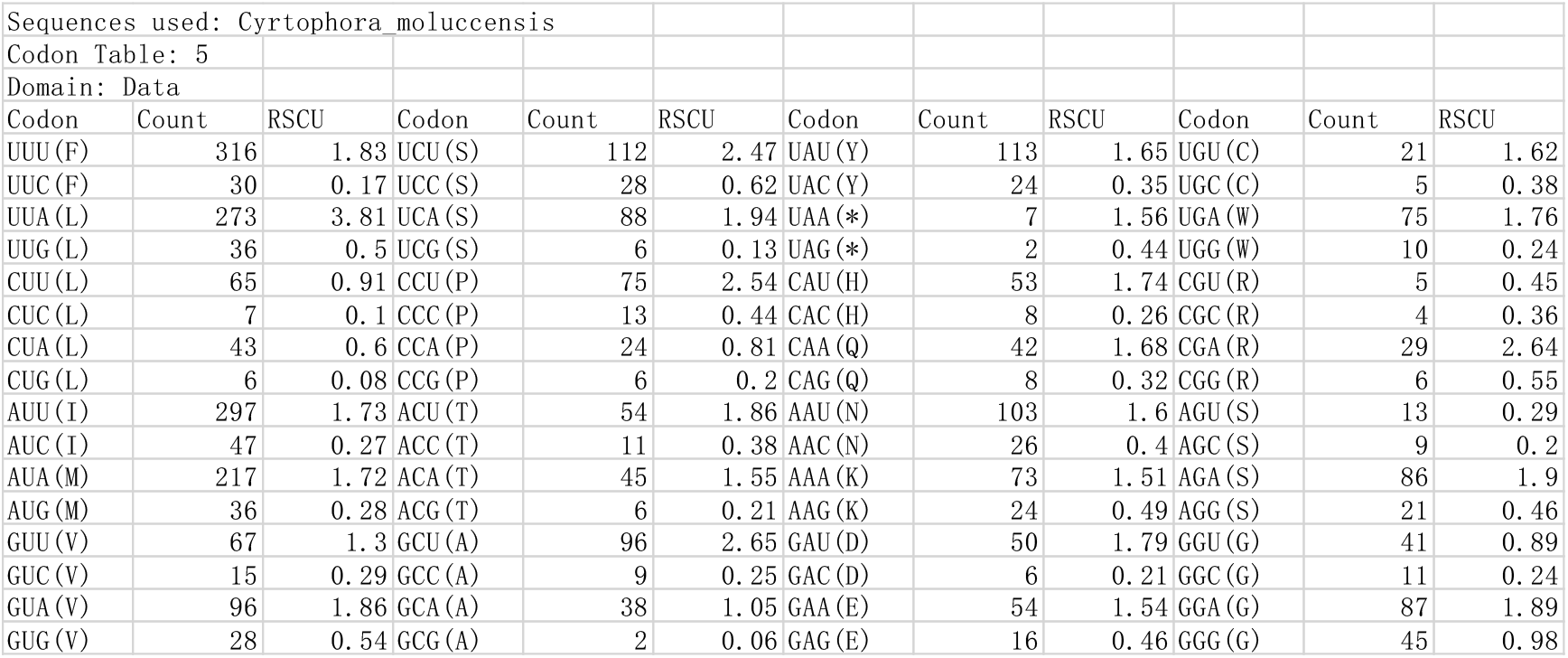
Relative synonymous codon usage (RSCU) table.

**Fig. S8.**
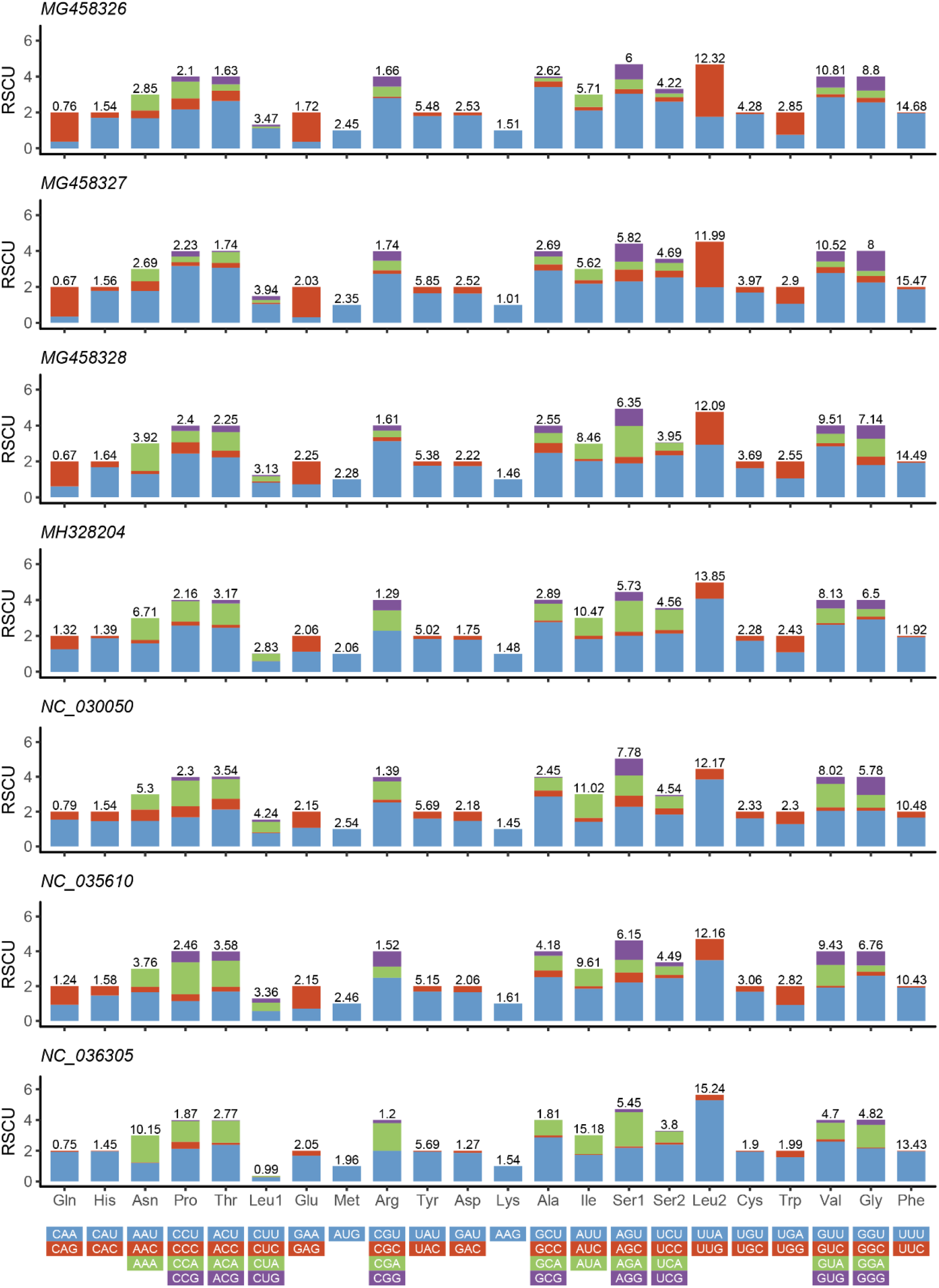
Relative synonymous codon usage (RSCU) of seven species.

**Fig. S9.**
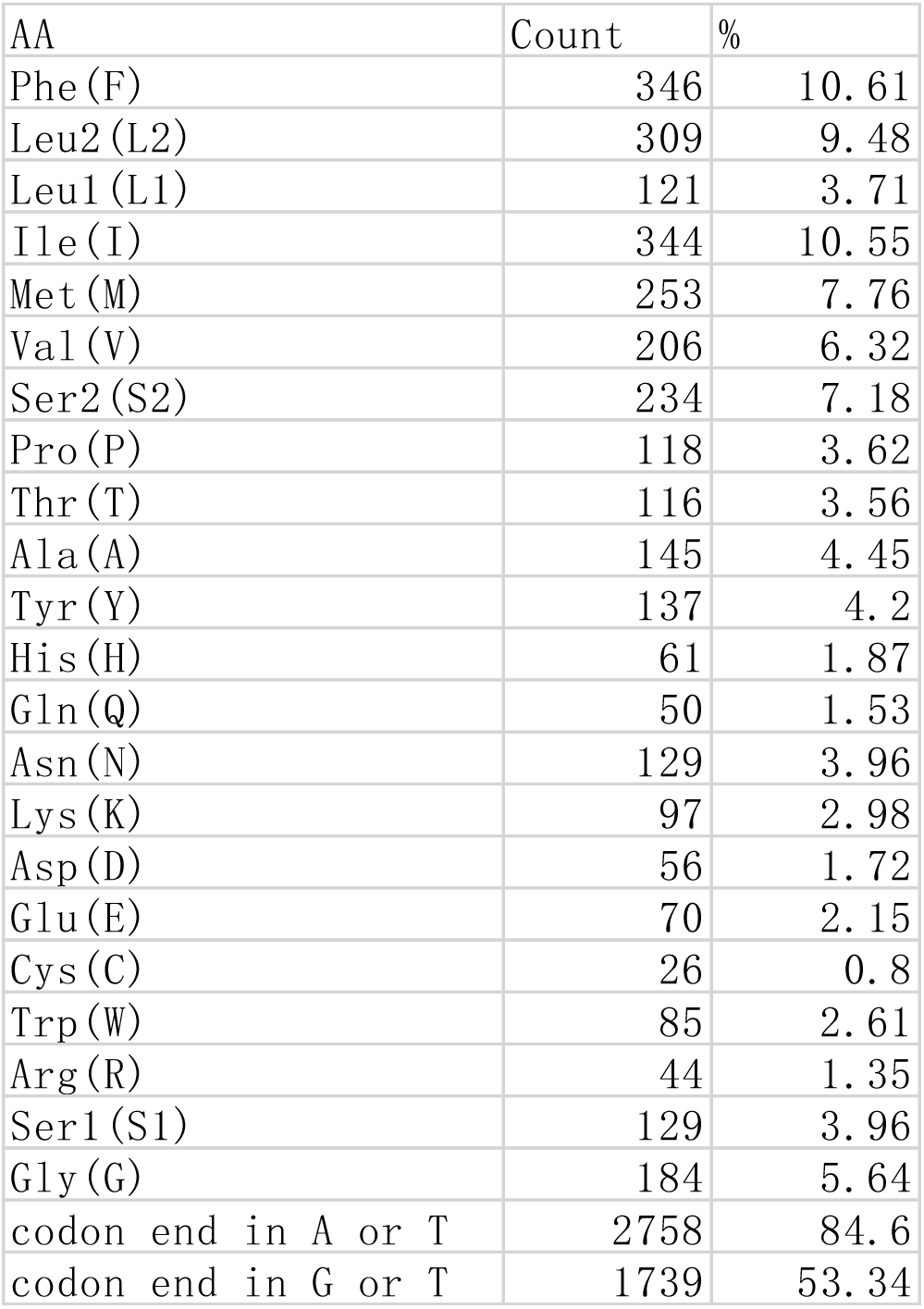
Amino acid usage statistics file.

**Fig. S10.**
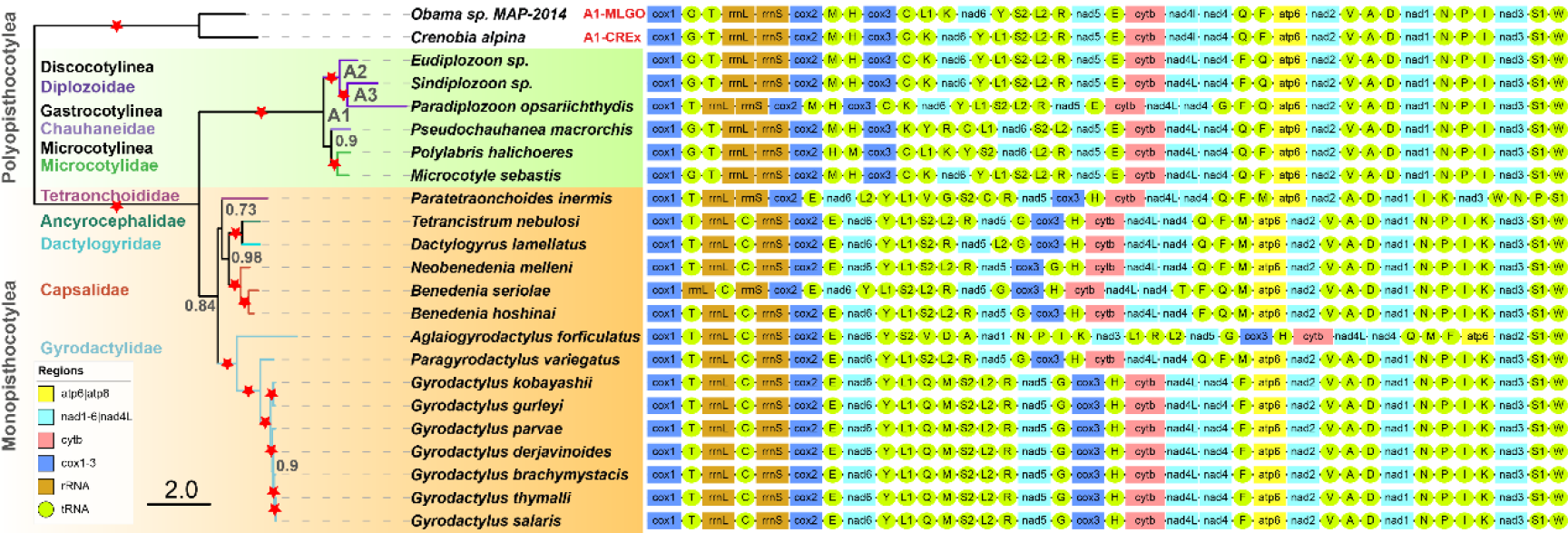
Mapping gene orders of monogenean mitogenomes onto the phylogenetic tree. The figure was published in Zhang, et al. (2018).

**Fig. S11.**
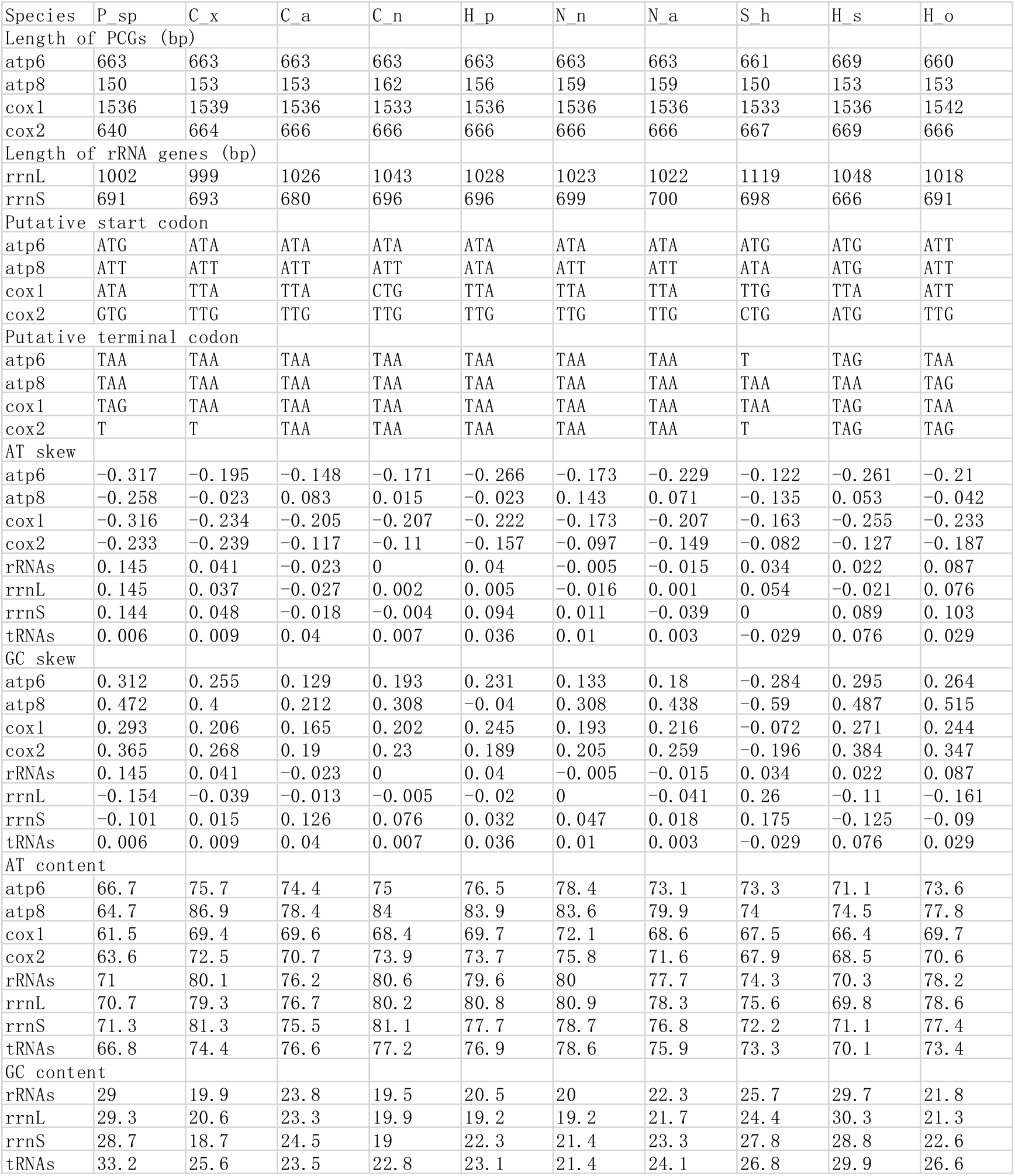
Statistics for individual genes, including size, start and terminal codons, base composition and skews. Only 4 protein-coding genes (PCGs) are shown.

**Fig. S12.**
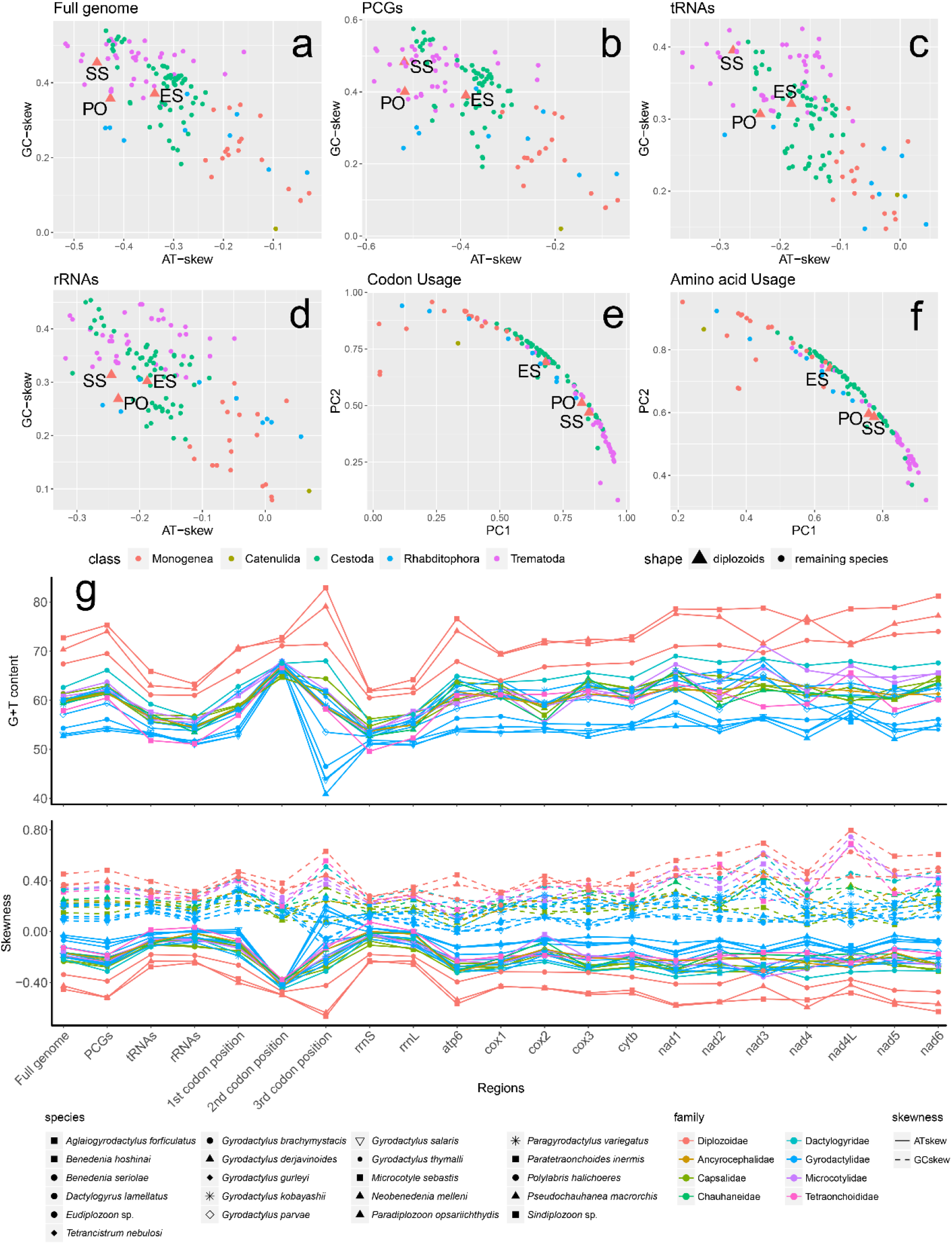
Skewness and base content of some flatworm mitogenomes. The figure was published in Zhang, et al. (2018).

**Fig. S13.**
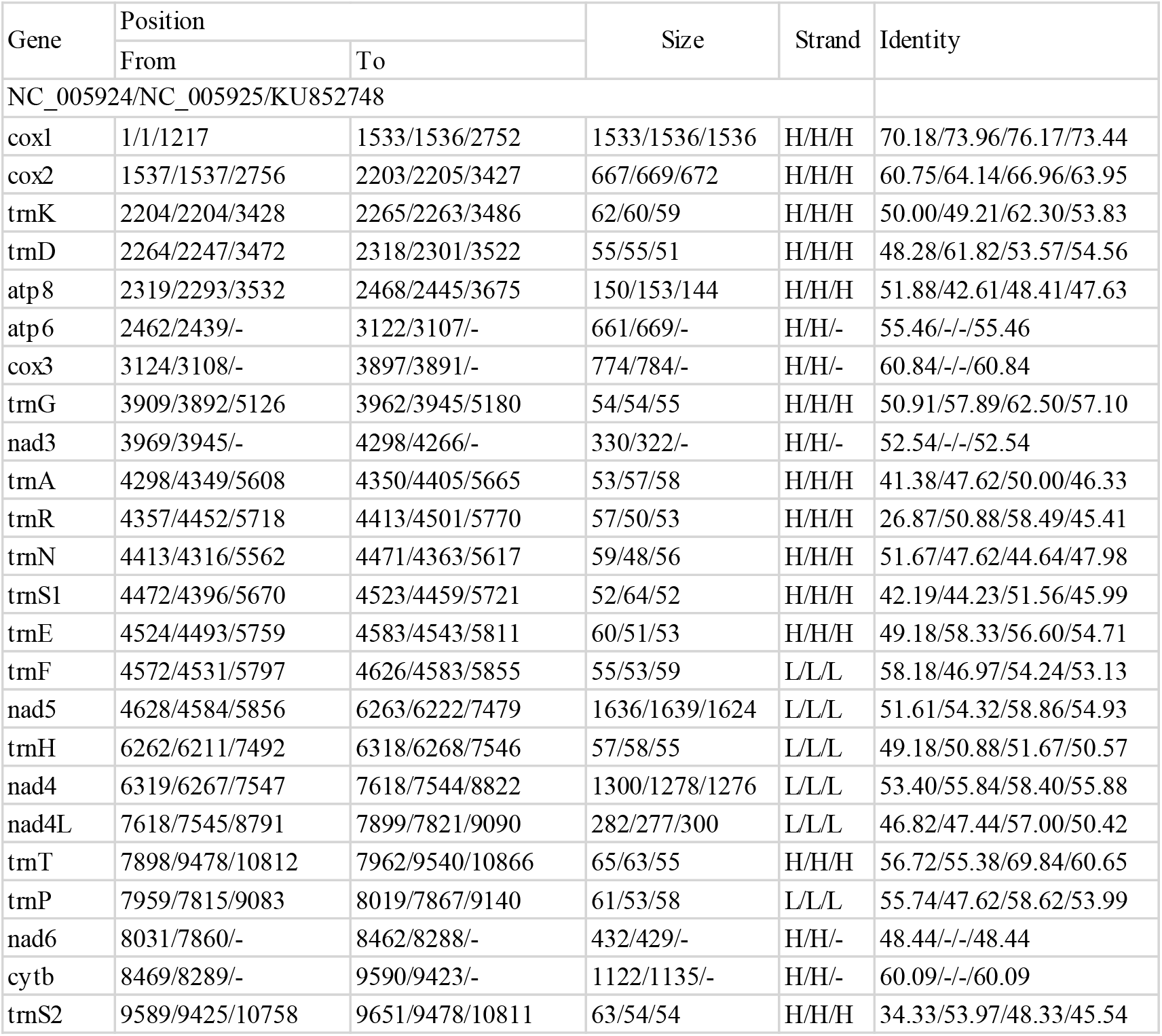
Comparison of genomic annotations and pairwise similarity calculation for homologous genes.

### Usage examples

PhyloSuite has been used previously to conduct analyses in a number of published papers. You may refer to the following publications for examples of the use of PhyloSuite: (Hua, et al., 2018;Li, et al., 2018; Li, et al., 2017; Liu, et al., 2017; Liu, et al., 2018; Liu, et al., 2018; Wang, et al., 2017; Wen, et al., 2017; Xi, et al., 2018; Zhang, et al., 2018; Zhang, et al., 2018; Zhang, et al., 2018; Zhang, et al., 2017; Zhang, et al., 2017; Zou, et al., 2017; Zou, et al., 2018). Note that we have merged our two older beta tools, MitoTool (https://github.com/dongzhang0725/MitoTool) and BioSuite (https://github.com/dongzhang0725/BioSuite) into PhyloSuite, so some of our older published papers may refer these two tools instead.

